# Diagnostic optical sequencing

**DOI:** 10.1101/685792

**Authors:** Lee E. Korshoj, Prashant Nagpal

## Abstract

Advances in precision medicine require high-throughput, inexpensive, point-of-care diagnostic methods with multi-omics capability for detecting a wide range of biomolecules and their molecular variants. Optical techniques have offered many promising advances towards such diagnostics. However, the inability to squeeze light with several hundred-nanometer wavelengths into angstrom-scale volume for single nucleotide measurements has hindered further progress. Recently, a block optical sequencing (BOS) method has been shown for determining relative nucleobase content in DNA k-mer blocks with Raman spectroscopy, and a block optical content scoring (BOCS) algorithm was developed for robust content-based genetic biomarker database searching. Here, we performed BOS measurements on positively-charged silver nanoparticles to achieve 93.3% accuracy for predicting nucleobase content in DNA k-mer blocks (where k=10), as well as measurements on RNA and chemically-modified nucleobases for extensions to transcriptomic and epigenetic studies. Our high-accuracy BOS measurements were then used with BOCS to correctly identify a β-lactamase gene from the MEGARes antibiotic resistance database and confirm the *Pseudomonas aeruginosa* pathogen of origin from <12 content measurements (<15% coverage) of the gene. These results prove the integration of BOS/BOCS as a diagnostic optical sequencing platform. With the versatile range of available plasmonic substrates offering simple data acquisition, varying resolution (single-molecule to ensemble), and multiplexing, this optical sequencing platform has potential as the rapid, cost-effective method needed for broad-spectrum biomarker detection.

## Introduction

Precision medicine can become one step closer to realization with inexpensive, non-specific assays capable of broad-spectrum diagnostics. Ideally, a single point-of-care test could rapidly screen an array of genomic, transcriptomic, and epigenetic biomarkers to inform proper therapeutic treatment.^1,2^ One urgent application is to combat antibiotic-resistant infections, which continue to create an increasing financial strain on healthcare systems worldwide and lead to high rates of mortality for the millions affected each year.^3,4^ Clinicians carry the burden to rapidly assess the potential resistances of pathogens and administer the correct antibiotics from the onset of infection. Unfortunately, initial antibiotic regimens are incorrect as much as 50% of the time.^5–7^ As initial administration of ineffective antibiotic treatments has proven to result in higher mortality rates, where in some cases survival decreases by 8% per hour, it becomes clear that more effective diagnostics are needed.^8–11^ Current resistance diagnostics and pathogen profiling are often only performed after initial antibiotics fail. Most of these assays rely on cell cultures, PCR amplifications, and microarray analyses.^12–19^ These tests require hours to days to complete and carry significant costs, while many are specific for merely one or a few well-characterized strains or resistances. Therefore, a method to rapidly and affordably screen a wide range of drug resistance in clinically relevant microbial pathogens is vital for prescribing patients with appropriate and timely treatments that reduce mortality rates as well as the spread of further resistance. Such a method would not be limited to antibiotic resistance screening, but also applicable in the screening for cancers and other genetic diseases where early detection is critical for patient survival.^20–22^

Here, we address the clinical diagnostic needs discussed above by demonstrating an optical sequencing platform capable of rapid, broad-spectrum diagnostics. Our diagnostic optical sequencing combines label-free Raman spectroscopy measurements of A-G-C-T (adenine-guanine-cytosine-thymine) content within DNA fragments (block optical sequencing – BOS^23^) with a robust bioinformatics algorithm for genetic biomarker database searching (block optical content scoring – BOCS^24^). BOS uses plasmonic substrates for surface-enhanced Raman spectroscopy (SERS) measurements of nucleotide content within unlabeled DNA k-mer blocks as an alternative to single-letter sequencing. Given the range of available plasmonic metallic nanoparticles and nanopatterned surfaces, BOS is flexible to many applicable substrates and can be made high-throughput with millions of simultaneous SERS measurements through multiplexing.^25–30^ BOCS uses a machine learning approach for mapping and scoring k-mer block reads from BOS to a genetic biomarker database. Initial simulations of BOCS showed accurate recognition of specific resistance, cancer, and other genetic disease genes with significantly less than full coverage of the genes.

We successfully coupled BOS measurements with the BOCS algorithm for characterization of a β-lactamase gene within the pathogen of origin. Specifically, we show that merely a few highly accurate BOS readings of DNA k-mer block content (<< full coverage of the gene) from silver nanoparticles can be used with the BOCS algorithm to identify the correct OXA β-lactamase (class D) gene from a comprehensive antibiotic resistance database and confirm the *Pseudomonas aeruginosa* pathogen from which it originates. While our initial BOS method employed a multiplexed, silver-coated nanopyramid substrate for SERS, we utilized metallic nanoparticles here to demonstrate broader applicability and varying (single-molecule versus ensemble). We also show extensions to transcriptomics and epigenomics. Ultimately, the results here demonstrate the use of combined BOS/BOCS as a diagnostic optical sequencing platform for inexpensive and rapid identification of broad-spectrum genetic, transcriptomic, and epigenomic biomarkers.

## Results and discussion

### BOS measurements with positively-charged silver nanoparticles

Due to the distinct biochemical nature of the nucleobases in DNA, each has unique interactions with light photons and a unique Raman spectrum.^31–34^ With SERS measurements, surface plasmon polaritons squeeze light photons with several hundred-nanometer wavelengths into molecular length scales, where they can interact with the nucleotides in DNA.^25,35,36^ Despite the advances in nano-optics and three-dimensional plasmonic nanofocusing, the nanometer scale mode volumes prevent characterization of photon interactions with single nucleotides. While this inhibits single-letter sequencing of DNA, it allows for determination of A-G-C-T content within k-mer blocks. By measuring content instead of single-letter sequences, which is more than sufficient for biomarker detection, the BOS system remedies the massive data analysis problem present with other sequencing platforms by incorporating inherent lossy data compression, and has the potential for extremely high-throughput measurements with millions of simultaneous multiplexed reads.^36–38^

In this study, we collected BOS measurements from ssDNA k-mer blocks with positively-charged, spermine-coated silver nanoparticles (Ag NPs) as the plasmonic substrate (Figure 1a). Recent work has shown strong, reproducible SERS signal from a range of substrates like single DNA molecules on nanopyramid substrates,^23^ and ensemble measurement from ∼25 nm cationic nanoparticles for ssDNA, dsDNA, and RNA.^29,39–41^ The Ag NPs remain stable in colloidal solution due to electrostatic repulsion of the positively-charged ligands, and show no significant background Raman signal (Figure S1). SERS signal is only achieved upon aggregation with the addition of negatively-charged nucleic acids, which is DNA k-mer blocks for our measurements. As seen from the extinction spectrum in Figure 1a, a strong localized surface plasmon resonance (LSPR) peak is observed at ∼392 nm for blank Ag NPs, and a large red shift is observed with the addition of DNA due to aggregation. When added to the Ag NPs, the DNA strands attach electrostatically to the nanoparticle surfaces and at interparticle hot spots leading to strong SERS excitation with a 532 nm Raman laser. In the analysis of all SERS measurements shown here, we perform a consistent signal processing (cosmic ray removal, smoothing, shift correction – Figure S2) and normalization (baseline subtraction, normalization to standard peak – Figure S3).

**Figure 1.**
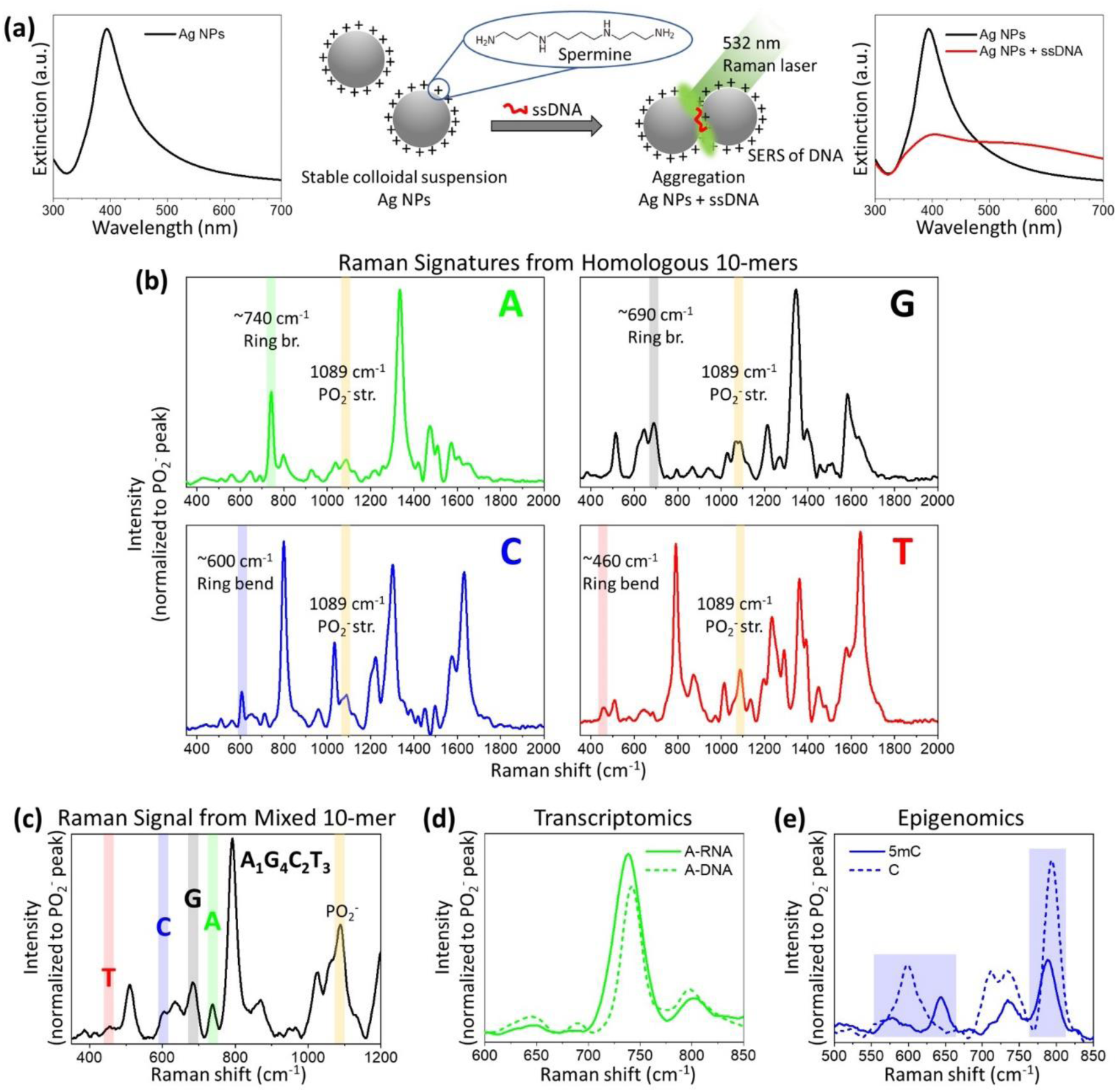
Overview of the BOS system with positively-charged silver nanoparticles, (a) SERS measurements of ssDNA k-mer blocks are collected from colloidal suspensions of positively-charged Ag NPs with a 532 nm laser. Signal enhancement is achieved via aggregation of the Ag NPs in the presence of negatively-charged DNA k-mer blocks, as evident by the red shift in the extinction spectrum, (b) Raman signatures for the four DNA nucleobases A, G, C, and T collected from homologous 10-mer sequences. The signatures provide the most distinctive Raman mode peaks for each base, which are used to deconvolute the content of mixed sequence k-mer blocks. These “signature peaks” are marked along with the 1089 cm ^−1^ PO_2_^−^ str. peak used for normalization (A: ∼740 cm^−1^ ring br., G: ∼690 cm ^−1^ ring br., C: ∼600 cm^−1^ ring bend, T: ∼460 cm^−1^ ring bend), (c) In mixed sequence DNA blocks, the four signature peaks are present with relative intensities (normalized to the PO_2_^−^ peak) corresponding to their respective content, (d) Raman signal for RNA and DNA show near identical shifts (shown for adenine, A, in RNA and DNA), demonstrating the potential for transcriptomic BOS analyses, (e) Subtle perturbations are seen in the Raman signal due to nucleobase chemical modifications (shown and highlighted for the cytosine, C, modification to 5-methylcytosine, 5mC), demonstrating the potential for epigenomic BOS studies.

For sequencing applications it is essential to first know the specific Raman signal, or signature, for each of the four nucleobases A, G, C, and T. To get these signatures, we performed SERS measurements on homologous 10-mer DNA sequences (i.e., poly(N)_10_, where N is A, G, C, and T). Shown in Figure 1b, there exists a complex pattern of Raman peak features for each nucleobase over the range of 350 – 2000 cm^−1^ shift (assignment of specific Raman peak modes provided in Table S1). For BOS, we identified a single distinctive Raman peak for each nucleobase as the “signature peak” which is later used to determine content in unknown sequence blocks. For purines, we selected the ring breathing modes at ∼740 cm^−1^ for A and ∼690 cm^−1^ for G.^29,31,34,40,42^ For pyrimidines, we selected the ring bending modes at ∼600 cm^−1^ for C and ∼460 cm^−1^ for T.^29,32,33,40,42^ Within each SERS measurement, the PO_2_-stretching mode peak at 1089 cm^−1^ due to the phosphate backbone is used as an internal standard for normalizing the relative peak intensities, as is consistent with other studies employing nanoparticle substrates.^27,40^ All signature peaks and the PO_2-_ normalization peak are highlighted in Figure 1b. In the SERS signal from a mixed sequence DNA block, the four signature peaks are present with relative intensities (normalized to the PO_2-_ peak) corresponding to their respective content, shown for a 10-mer DNA block with content A:1, G:4, C:2, and T:3 in Figure 1c. These relative intensities in signature peak locations can therefore be used to deconvolute the signal from an unknown mixed sequence.

While the focus in this study is put on DNA, it is important to note that impactful extensions exist for transcriptomics and epigenomics by applying BOS to RNA and chemically modified nucleobases. As shown in Figure 1d for a homologous RNA sequence of repeating adenine, A, the signature peaks are nearly identical between RNA and DNA. This is evident by the similar ∼740 cm^−1^ ring breathing mode peak for A, which is consistent with recent work by other groups.^41^ Additionally, small perturbations to the Raman spectrum can be seen due to modified nucleobases like the modification of cytosine, C, to 5-methylcytosine, 5mC (also measured from homologous sequences), which plays an important role in gene regulation.^43,44^ Seen in the highlighted regions of Figure 1e, we observe shifts and intensity changes to the signature peak of C at ∼600cm^−1^ and the strong ring breathing mode for pyrimidines at ∼800 cm^−1^, which are consistent with previous studies.^26^ This opens the opportunity to directly apply our diagnostic optical sequencing platform to transcriptomic and epigenomic studies.

### Calibrating the BOS system with DNA block standards

To fully deconvolute the A-G-C-T content of an unknown mixed sequence DNA k-mer block, it is necessary to know the full range of intensity values for signature peaks of each nucleobase. Therefore, we used custom DNA k-mer blocks with known content as standards for generating content calibrations for BOS. The 14 calibration blocks are provided in Table 1. These 14 ssDNA 10-mer calibration blocks span the range of 0 – 1 fractional content for each of the four nucleobases. Blocks Cal_1, Cal_2, Cal_3, and Cal_4 provided the Raman signatures shown in Figure 1b, as they are of content = 1. Together, SERS measurements on the set of 14 calibration blocks were used to generate the BOS calibrations shown in Figure 2 (all SERS spectra of the calibration blocks are provided in Figure S4). On the left of Figure 2, SERS spectra are plotted for increasing fractional content of a particular nucleobase (lighter to darker shades) in the zoomed-in region of the signature peaks. A direct, linear correlation is observed between fractional content and the signature peak normalized intensity for each of the nucleobases, seen in the fitted data points on the right of Figure 2 (data points and variance are from five technical replicates of each calibration block). The trends are all linear although the range of measured intensity values can vary significantly. Intensities for the adenine, A, signature peak at ∼740 cm^−1^ range from 0 to >4 while the intensities for the thymine, T, signature peak at ∼460 cm^−1^ range from 0 to 0.3. Linear fits to these trends, with intercept locked at zero, provide the finalized correlations which can be used to determine content in unknown DNA k-mer blocks.

**Table 1.**
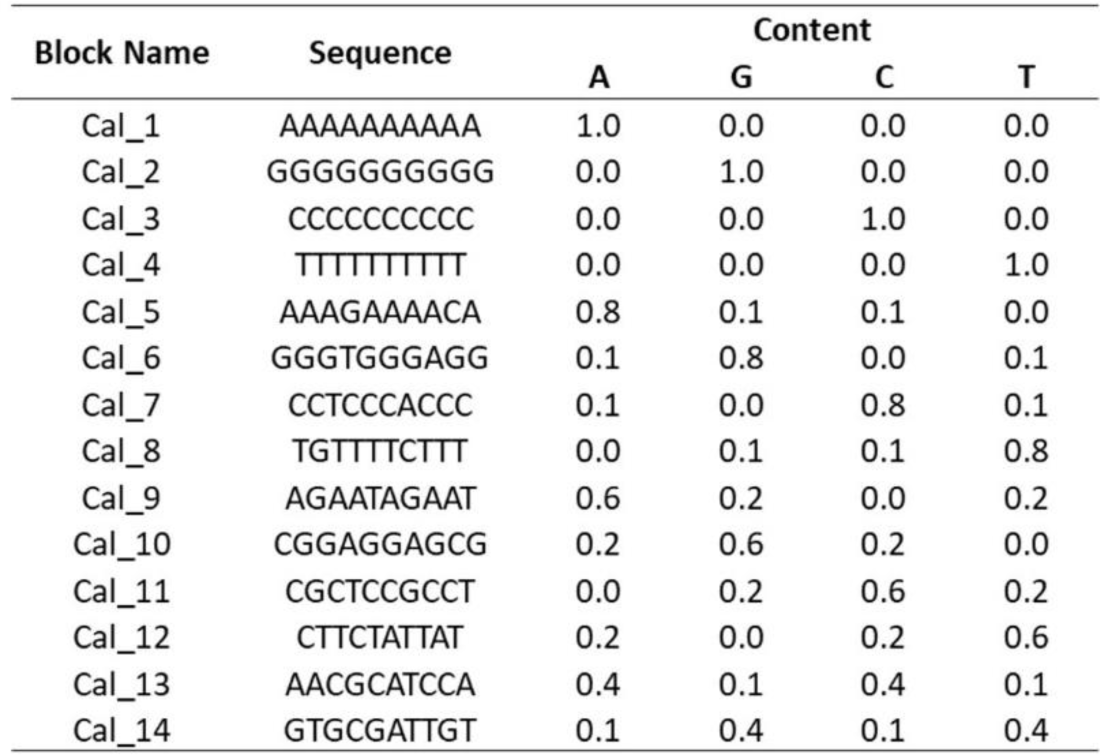
calibration blocks (ss DNA 10-mers)

**Figure 2.**
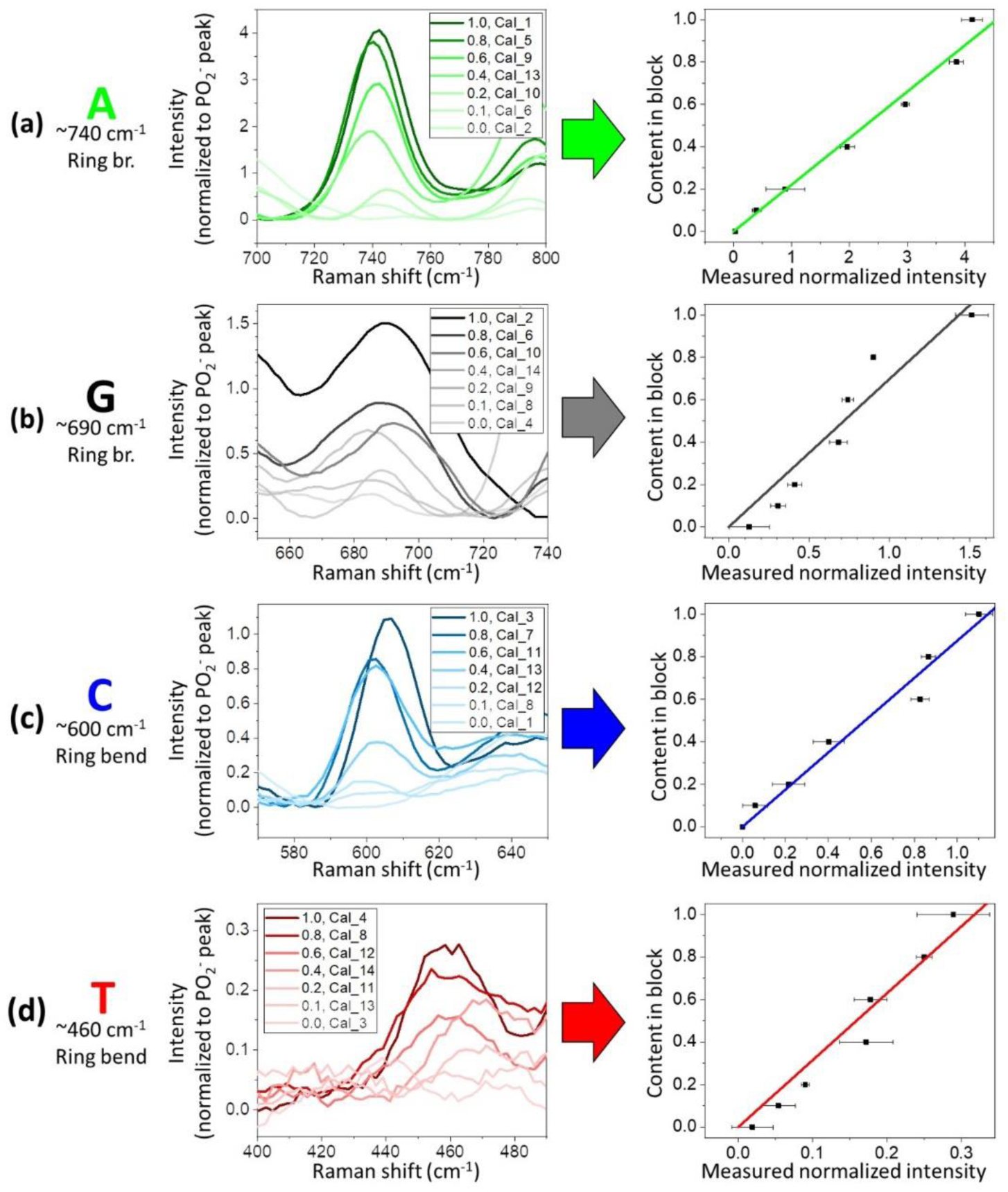
BOS calibrations. Analyzing the correlations between varying nucleobase content within the DNA 10-mer calibration blocks from Table 1 and changes in the signature peak intensity for (a) A: ∼740 cm ^−1^ ring br., (b) G: ∼690 cm ^−1^ ring br., (c) C: ∼600 cm^−1^ ring bend, and (d) T: ∼460 cm^−1^ ring bend. (Left) For each nucleobase, increasing content within a block (lighter to darker shades, labeled in the plot legend with the corresponding calibration 10-mer block from Table 1) leads to a linear increase in the intensity of the signature peak. (Right) Linear fits, with intercept locked at zero, of measured signature peak normalized intensity versus content within the block (data points and variance are from five technical replicates of each calibration block). These fits are used as calibrations to identify content in unknown mixed sequence DNA k-mer blocks.

### Content identification within gene blocks

We applied the BOS calibrations towards identifying content within k-mer blocks from an actual gene sequence, for subsequent integration with the BOCS algorithm. The 15 gene blocks are provided in Table 2. These 15 ssDNA 10-mer gene blocks are from an OXA β-lactamase (class D) gene found in *P. aeruginosa* (full sequence and mapped blocks are provided in Figure S5). From SERS measurements on the 15 gene blocks (all SERS spectra of the gene blocks are provided in Figure S6), we measured the normalized intensity for signature peaks (averaged from three technical replicates). Figure 3 details the process of using these measurements for predicting content within the blocks. Using the Gen_4 block as an example, the measured normalized intensity for each of the four signature peaks is used to predict a content for each nucleobase A, G, C, and T by multiplying the measured intensity by the slope of the linear calibrations. This initial raw predicted content is then normalized such that the sum is equal to one. Knowing that each nucleobase must be present in integer quantities, the final predicted content can be determined via rounding. The final predicted content for the Gen_4 block agrees with the actual content given in Table 2.

**Table 2.**
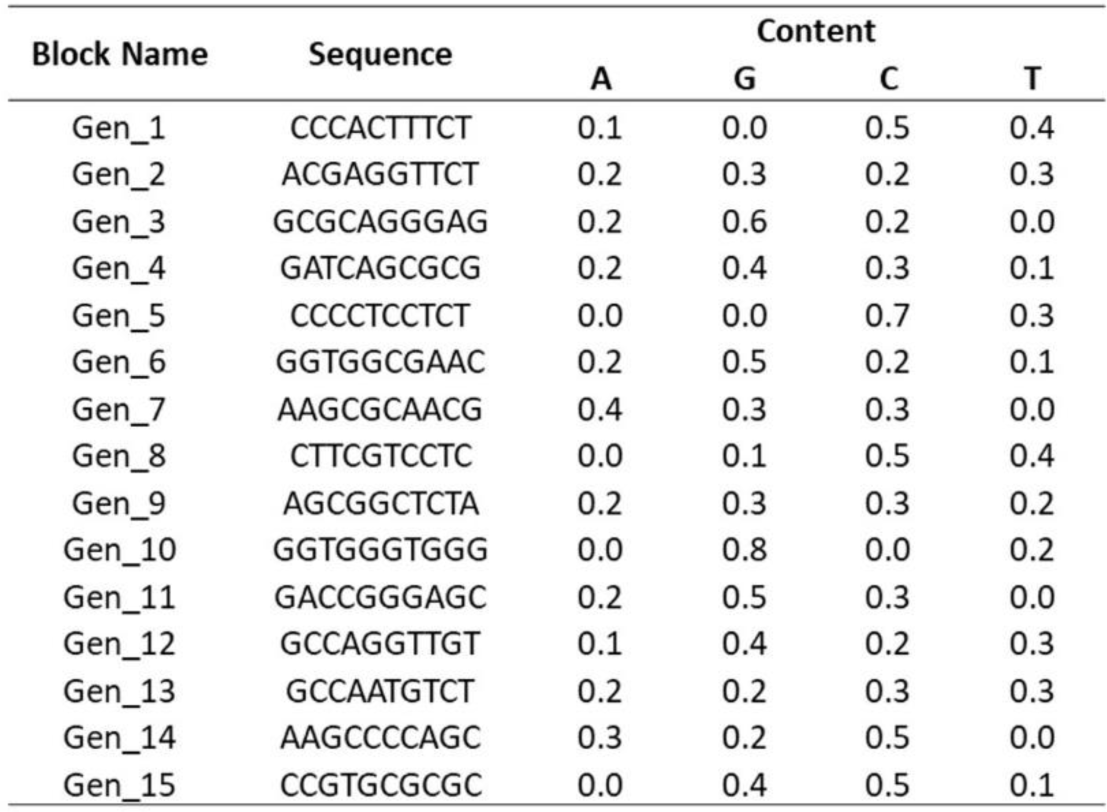
Gene blocks (ssDNA 10-mers)

**Figure 3.**
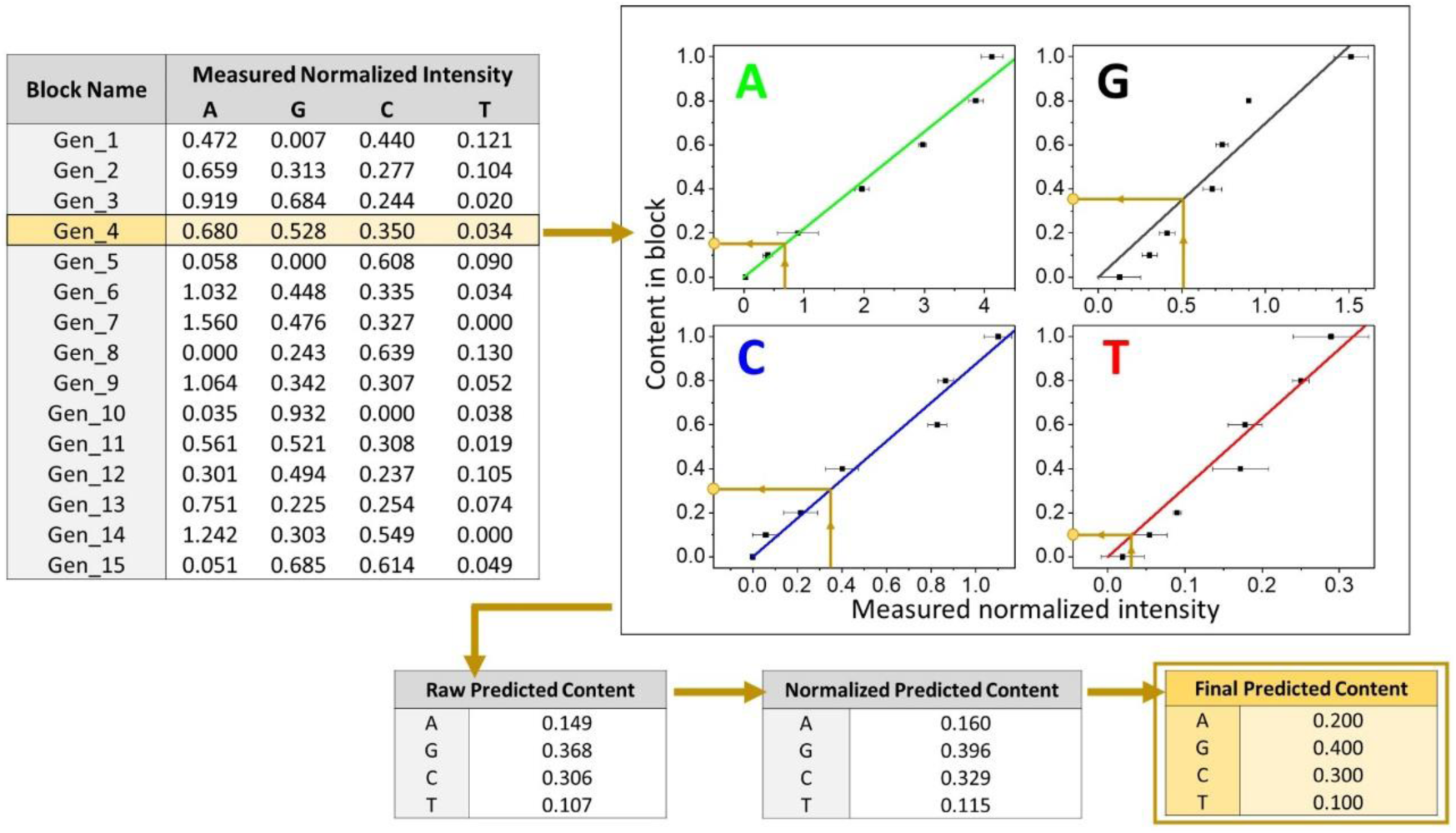
Content identification within gene blocks. The content of unknown mixed sequence DNA blocks (shown for the 15 10-mer gene blocks from Table 2) can be identified from the calibrations for each of the four nucleobases. Using block Gen_4 as an example, the measured normalized intensity for each of the four signature peaks (averaged from three technical replicates) is used to predict raw content. This raw content is then normalized such that the total predicted content equates to one. The normalized content is then rounded so that each base has an integer number within the block.

Predicted content for all 15 gene blocks is provided in Figure 4, where the predictions are compared to the actual content. It is important to note that the accuracies reported in BOS are different than those from traditional single-letter sequencing. Since reads are of A-G-C-T content and not letter-by-letter sequences, one misidentification results in a double error. This is because the content of the incorrect nucleobase and substituted nucleobase are both affected. A confusion matrix analysis on the single nucleobase level throughout all 15 blocks shows that the majority of errors result from guanine, G, content being under identified (∼10% of G bases throughout the gene blocks). In total, errors in the predicted content were present in only four of the blocks – three which resulted from a single nucleobase swap (80% accuracy for the block) and one which double-swapped nucleobases (60% accuracy for the block). Content in the other 11 blocks was predicted with 100% accuracy. Overall, the content was predicted at an average accuracy of 93.3%.

**Figure 4.**
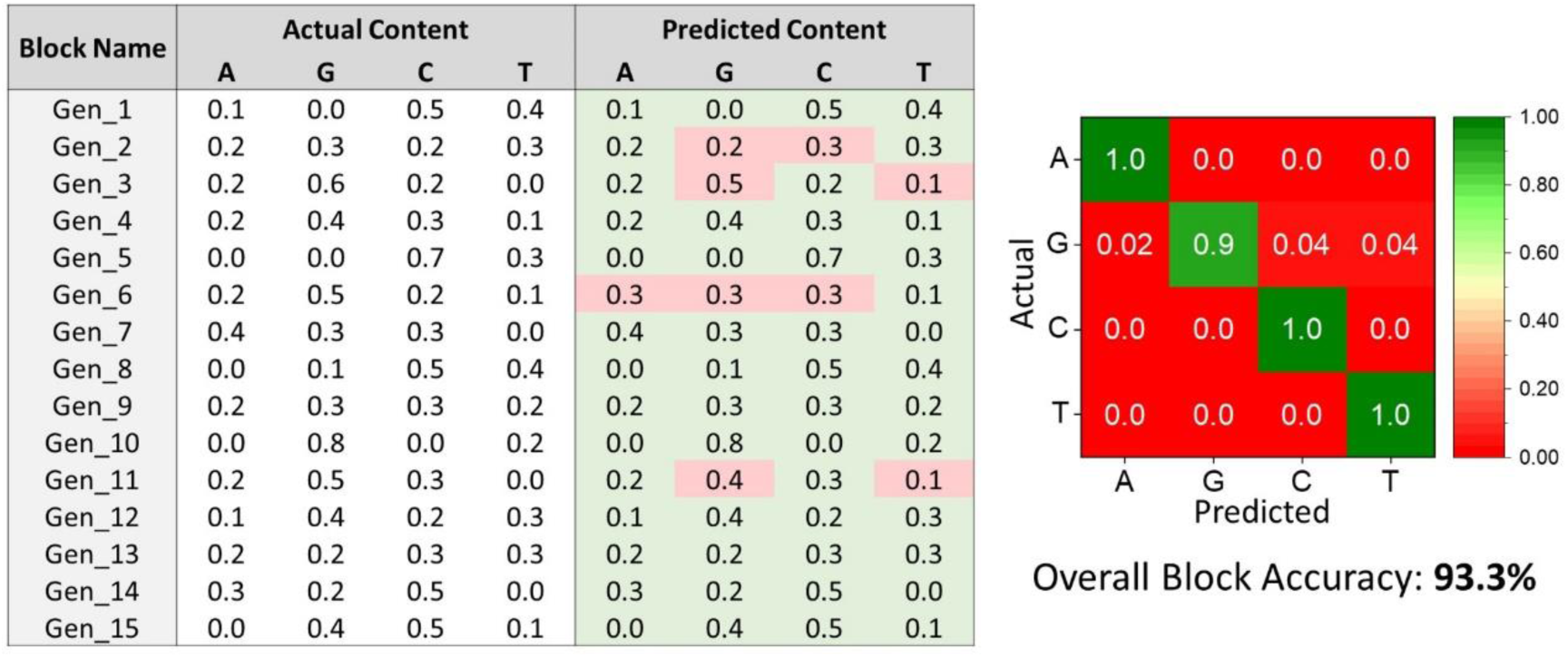
Highly accurate content identification. Actual and predicted content is compared for the 15 10-mer gene blocks from Table 2. Since BOS relies on content and not letter-by-letter sequences, one misidentification results in a double error because the content of the incorrect nucleobase and substituted nucleobase are both affected. In the figure table, correct predictions are highlighted in green, and incorrect predictions are highlighted in red. A confusion matrix analysis on the single nucleobase level shows that the majority of errors result from guanine, G, content being under identified (∼10% of G bases throughout the gene blocks). In total, the content for the 15 gene blocks was identified at an average accuracy of 93.3%.

### MDR pathogen profiling with BOS and BOCS

For full integration into a diagnostic method, the high-accuracy BOS reads were coupled with the BOCS algorithm for genetic biomarker detection. With the combined BOS/BOCS platform, we set out to demonstrate detection of a *P. aeruginosa* infection with drug-resistant β-lactamase gene. *P. aeruginosa* is a clinical multidrug-resistant (MDR) pathogen of critical importance due to its prevalence for causing bloodstream, urinary, and pulmonary infections in hospital settings, especially for immunocompromised patients in intensive care settings.^45,46^ Due to the multiple mechanisms of inherent and acquired resistance of this organism, patients infected with *P. aeruginosa* have limited therapeutic options. It is therefore imperative to have more early-stage, rapid diagnostic techniques in place to screen for *P. aeruginosa* so that effective antibiotic regimens can be prescribed from the onset of infection.

The BOCS algorithm was developed to specifically utilize DNA k-mer block content reads from the BOS system.^24^ It operates analogously to probability-based sequence analyzers such as those employed for peptide identification from mass spectrometry data,^47–49^ and alignment programs used for mapping next-generation sequencing reads to reference genomes.^50,51^ In a similar fashion, the BOCS algorithm relies on probabilistic content alignments to database sequences of genetic biomarkers. Outlined in Figure 5a, the algorithm cycles through each of the logged k-mer block content reads from BOS and performs a content-based alignment with each gene sequence in a database, translating through the gene sequence one nucleotide at a time. The alignment tracks the number of match locations – where the k-mer block content matches the content of the k-length gene segment. The number of match locations is the fundamental parameter in a set of six probability factors that act as machine learning elements in the calculation of an overall content score. Genes in the database are probabilistically ranked, and identified, based on the content score as it compounds with more blocks being analyzed. The algorithm also incorporates logic elements such as a penalty score given to genes in the database where no matches are found during alignment, thresholding to eliminate low-ranking genes in the database that may skew the content scoring, and entropy screening to eliminate BOS reads that have a maximal number of permutations based on the content. Thorough simulations of BOCS with antibiotic resistance, cancer, and other genetic disease databases proved very robust, even under the pressures of variable k-mer block lengths, high error rates, and in the presence of blocks comprised of multiple genes.

**Figure 5.**
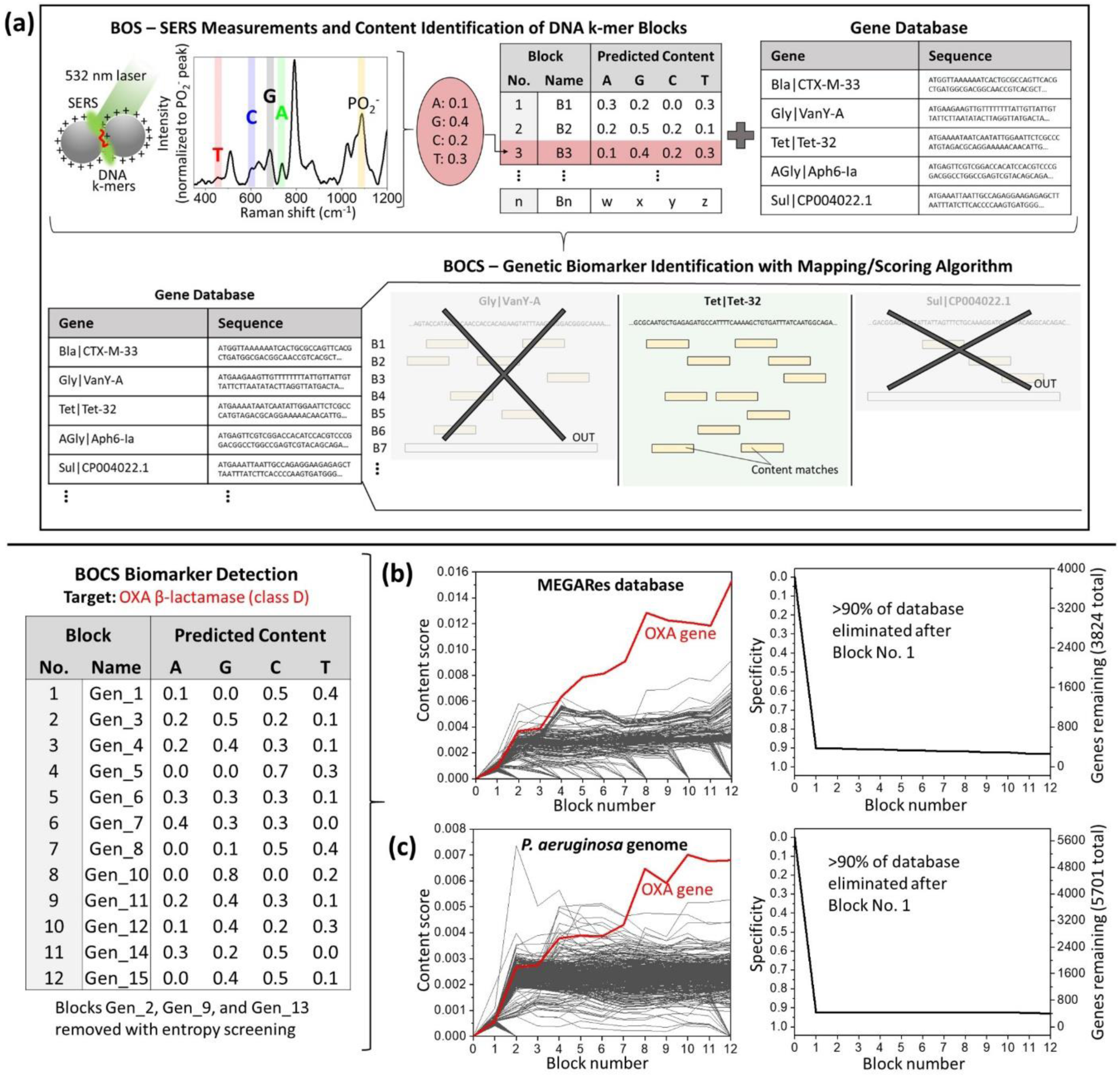
MDR pathogen profiling with integrated BOS and BOCS. (a) Overview of the BOCS algorithm integration with BOS measurements. Starting with a log of measured content within DNA k-mer blocks from BOS (B1…Bn as shown) and a gene database (excerpts from the MEGARes antibiotic resistance database are shown), the blocks are individually aligned to each gene in the database based on content. This alignment consists of finding all match locations for the k-mer block content within a gene via translating through the gene one nucleotide at a time and looking at fragments of length k. For each block, a content score is calculated based on the number of matches for the k-mer block and various probability factors. As more blocks are analyzed, content scores are compounded and genes in the database are ranked and eliminated. The BOCS algorithm was run for the 15 10-mer gene blocks in Table 2 from an OXA |3-lactamase gene (with the predicted content at 93.3% accuracy). Note that only 12 of the 15 blocks were used, as three were eliminated with entropy screening. Two cases were studied: 1. Identifying the gene from the MEGARes antibiotic resistance database of ∼4000 resistance genes (b), and 2. Identifying the gene within the *P. aeruginosa* genome (c). Both cases demonstrate robust identification of the correct OXA resistance gene from content score ranking, requiring merely a few content measurements. Additionally, >90% of genes in both databases were eliminated after a single block was analyzed by the BOCS algorithm. The following settings were used when running BOCS: penalty score - 0.1, thresholding multiplier-0.1, entropy screening - ‘on’ (eliminated only the blocks with permutations > 25000).

We ran our measured gene blocks with predicted content at 93.3% accuracy through the BOCS algorithm against the MEGARes antibiotic resistance database^52^ comprised of ∼4000 known resistance genes, including the OXA β-lactamase (class D) gene of our measured gene blocks. This analysis demonstrates the ability of the BOS/BOCS platform to diagnose antibiotic resistances from unknown samples with no prior knowledge of pathogen or strain. The table of gene blocks and their predicted content which was provided to the BOCS algorithm is shown in the lower portion of Figure 5. Note that three of the blocks were eliminated with the entropy screening functionality of BOCS because these blocks had predicted content with a maximum number of permutations/entropy (>25000 permutations of the 2-2-3-3 content within the 10-mer block), and therefore do not benefit the scoring and ranking. Figure 5b plots the content score as consecutive blocks have been analyzed (1 through 12 total blocks). It can be seen that the correct OXA gene from the MEGARes database is identified based on its top content score ranking well within the 12 blocks that are shown. The OXA gene begins to separate itself in ranking from the other database genes after merely the fifth block was analyzed, and it becomes easily identifiable after the twelfth block. The fifth block corresponds to only 0.063 or 6.3% coverage of the gene (i.e., 50 nucleotides analyzed in the five 10-mer blocks, out of 789 total nucleotides in the resistance gene sequence), while the twelfth block corresponds to 0.152 or 15.2% coverage of the gene. Also shown in Figure 5b is the specificity, or how significantly the gene database can be narrowed as consecutive blocks have been analyzed. We see that >90% of the total MEGARes genes in the database can be eliminated after merely the first block is analyzed.

Extending diagnostic applications further, we ran our measured gene blocks through the BOCS algorithm again after substituting the MEGARes database for the *P. aeruginosa* reference genome PAO1^53^ containing the OXA β-lactamase (class D) gene. This analysis indicates the ability of the BOS/BOCS platform to confirm pathogens and specific strains responsible for the infection. It also shows the robustness of the BOCS algorithm in identifying specific genes in the background of an entire microbial genome. Figure 5c plots the content score from BOCS as consecutive gene blocks have been analyzed. Just as with the MEGARes resistance database, the correct OXA gene from the *P. aeruginosa* genome database is identified based on its top content score ranking within the 12 blocks. Additionally, high specificity is seen as >90% of the total genes in the database can be eliminated after merely the first block is analyzed. These results demonstrate the potential for using our BOS/BOCS platform as a diagnostic optical sequencing technique for profiling MDR pathogens. The results shown here for a single β-lactamase gene in *P. aeruginosa* can be extended to other resistance genes from pathogenic microbial strains without any changes in experimental setup, ultimately providing the broad-spectrum detection needed for directing appropriate and timely treatments in a clinical setting.

## Conclusion

Sequencing methods like those from Illumina and Oxford Nanopore Technologies are transforming clinical diagnostics through rapid, cost-effective characterization of patient samples.^54^ For antibiotic resistance screening, current steps for pathogen characterization continue to rely on culture-based assays that can take days to weeks for results to be finalized. Next-generation sequencing has proven to significantly improve these metrics. The majority of next-generation techniques sequence a fragmented DNA sample and then piece together the fragments with alignment algorithms and a reference sequence.^37,51^ Clinical applications of Illumina sequencing suggest its use as an effective method of regular infection screening for control of MDR pathogens in hospitals.^55^ For this, 75-100× gene coverage is routinely needed for accurate results, which can require several days of runtime and analysis. The MinION system from Oxford Nanopore Technologies has shown the ability to identify antibiotic resistance genes and classify pathogens down to the species level.^56–58^ With these studies, runtimes were 6-18 hours and gene coverage was >10-14×, with optimized experimental protocols. Despite the progress being made with these sequencing methods, limitations of the platforms in both the measurements and algorithmic analysis continue to necessitate high coverages for accurate results, leading to long runtimes and higher costs.

Results from our BOS/BOCS platform demonstrate the significant potential of an optical sequencing technique to improve on the limitations seen in these current state-of-the-art sequencing methods. An optical technique could lead to improvements in time, cost, and complexity. On the measurement side, the label-free optical detection within BOS simplifies the sample preparation and gives the potential for multiplexing data acquisition, which would dramatically decrease runtimes. On the analysis side, the BOCS algorithm exceeds the performance of the alignment algorithms used by next-generation sequencers by requiring lower coverages. We showed that << 1 gene coverage (i.e., not even seeing the entire gene) is often all that is required for identification of genetic biomarkers from a database. This low coverage cuts down on costs associated with reagents as amplification of the sample could be significantly decreased or eliminated, and runtimes reduced.

In conclusion, we have coupled Raman measurements of A-G-C-T content in DNA k-mer blocks with a genetic biomarker database searching algorithm for integration as a diagnostic optical sequencing platform. With block optical sequencing (BOS), we employed positively-charged silver nanoparticles for reproducible SERS measurements from DNA blocks. Identification of four Raman ring breathing and bending modes, each specific to one of the four nucleobases, were observed to have intensities that are linearly correlated with increasing nucleobase content within a DNA block. A set of standard blocks with known nucleobase content were used to fit linear calibrations that led to 93.3% accuracy in predicting the relative A-G-C-T content of a set of blocks from a β-lactamase gene. With the block optical content scoring (BOCS) algorithm, we used our gene block content measurements to correctly identify β-lactamase resistance in the *P. aeruginosa* pathogen. Early detection of this opportunistic, nosocomial pathogen and its resistance to β-lactam antibiotics is crucial for providing patients, who are often immunocompromised, with effective antibiotic regimens. The powerful BOCS algorithm detected the specific OXA β-lactamase (class D) gene from the MEGARes antibiotic resistance database of ∼4000 genes without prior knowledge of pathogen or strain, and at merely <15% coverage of the gene. This can dramatically improve diagnosis times and costs compared to the culture-based detection strategies currently in place. Additionally, the nearly identical SERS signals seen between RNA and DNA provide evidence for directly translating the BOS/BOCS platform to the transcriptome. And detectable SERS variations due to chemically modified nucleobases such as 5-methylcytosine indicate the potential for epigenomic analyses. These results have demonstrated the use of the BOS/BOCS platform as a diagnostic optical sequencing technique, with extensions in genetics, transcriptomics, and epigenomics. Given the versatility of simple optical measurements and applicability to a wide range of biomarker databases, this platform has the potential for truly rapid and inexpensive, broad-spectrum diagnostics for personalized medicine.

## Experimental

### Materials

All custom DNA oligomers (10-mer calibration and gene blocks, Tables 1 and 2) and the custom homologous RNA oligomer (7-mer poly(rA)_7_) were ordered from Integrated DNA Technologies, Inc. (IDT). The custom homologous 5-methylcytosine oligomer (5-mer poly(5mC)_5_) was ordered from Gene Link. All oligomers were stored at −20°C or −80°C prior to use. Spermine tetrahydrochloride (99%), polyethyleneimine (PEI, branched, M.W. 10,000, 99%), and silver nitrate (99.9+%) were purchased from Alfa Aesar, and sodium borohydride from Fisher Chemical. All water used for synthesis, dilutions, and SERS measurements was ultrapure deionized (DI) water (Barnstead Thermolyne NANOpure Diamond purification system, water resistivity > 18 MΩ-cm).

### Ag NP synthesis

The synthesis protocol was adapted from van Lierop et al.^39^ Prior to synthesis, all glass vials were left to soak in PEI solution (0.4% v/v) overnight followed by extensive rinsing with ultrapure DI water. For Ag NPs, 40 µL silver nitrate solution (0.5 M) and 14 µL spermine tetrahydrochloride solution (0.1 M) were mixed with 20 mL ultrapure DI water and stirred for 20-30 min in dark. After 20-30 min, 500 µL of sodium borohydride solution (0.01 M) was spiked into the mixture (with continued stirring for 5-10 min). Ag NP colloids were allowed to sit overnight in dark (at room temperature), and the sediment at the bottom of the vial was then discarded.

### Sample preparation

Samples for SERS measurements were prepared by mixing 5 µL of DNA oligomer solution (100 µM in ultrapure DI water) with 495 µL of Ag NP colloidal solution, for a final DNA concentration of ∼1 µM. The DNA-Ag NP mixture was allowed to equilibrate for 2-3 h, followed by a quick sonication prior to measuring.

### UV-vis characterization

Ag NPs and their interaction with DNA were characterized with a VWR UV-1600PC spectrophotometer using 1 cm path length glass cuvettes. An LSPR peak at ∼392 nm was observed in the extinction spectrum for the blank Ag NP colloidal solution, with significant red shift seen upon the addition of DNA (Figure 1a).

### SERS measurements

SERS measurements were collected with a 532 nm 5 mW laser (Changchun New Industries Optoelectronics Tech Co., Ltd.) focused on the colloidal sample (500 µL volume) through an Olympus BX51 microscope with 5X 0.10 NA objective, and spectra were collected with a Princeton Instruments imaging spectrometer. For the calibration and gene blocks, five and three technical replicates were measured respectively under the same experimental conditions (30 s exposure time, 10 accumulations).

### SERS signal processing

Raw SERS spectra were processed by (1) Removing cosmic ray spikes in the spectrum, (2) 12-point average smoothing (±6 in each left/right direction), (3) Correcting small inconsistencies in the 1089 cm^−1^ PO_2-_ normalization peak. See Figure S2 for more details.

### SERS signal normalization

Processed SERS spectra were normalized by (1) Finding all baseline points along the spectrum between which linear fits are constructed, (2) Subtracting the linear fits between the baseline points, (3) Normalizing by the 1089 cm^−1^ PO_2-_ peak intensity. See Figure S3 for more details.

### BOCS algorithm

The BOCS algorithm, with details on the mapping and scoring aspects, was extensively described previously by Korshoj and Nagpal.^24^ The full code and databases to run simulations of BOCS can be found online at https://github.com/lkorshoj/Block-Optical-Content-Scoring.

## Acknowledgments

This work was supported by W.M. Keck Foundation, and partial support through National Science Foundation Soft Materials MRSEC at the University of Colorado through NSF Award DMR 1420736. L.E.K. acknowledges financial support from National Science Foundation Graduate Research Fellowship Program under Grant No. DGE 1650115.

