## Supplementary for "Diagnostic optical sequencing"

**Content:**

Figure S1. Background Raman signal from blank Ag NPs
Figure S2. SERS signal processing
Figure S3. SERS signal normalization
Table S1. Raman peak mode assignments
Figure S4. All SERS measurements from calibration blocks
Figure S5. OXA β-lactamase (class D) gene and blocks
Figure S6. All SERS measurements from OXA gene blocks

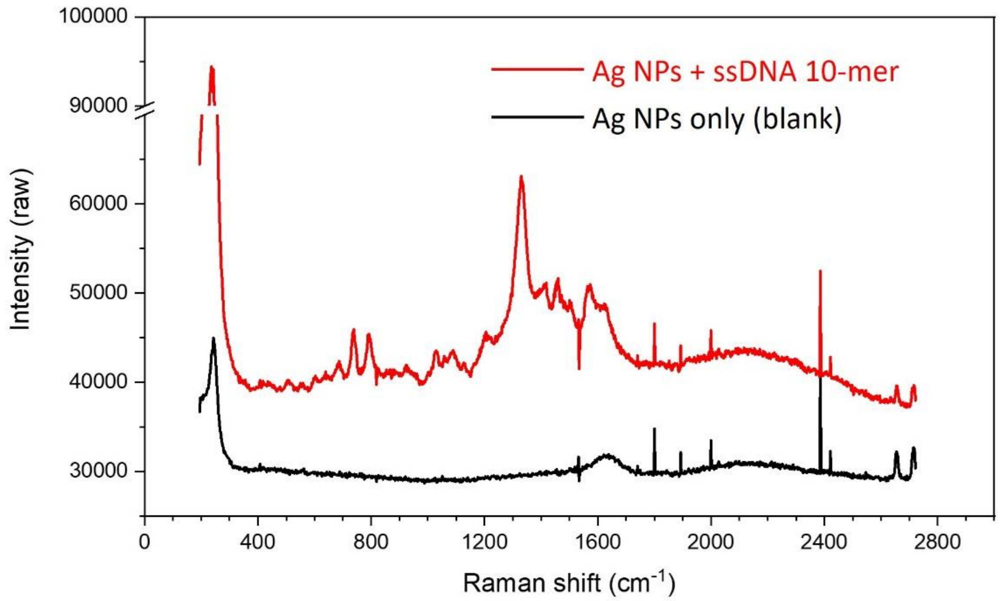


**Figure S1.** Background Raman signal from blank Ag NPs. Blank colloidal solutions of Ag NPs in the absence of DNA (black) show negligible Raman signal, especially in the 400-800 $cm^{-1}$ region where characteristic nucleotide signature peaks are found. Ag NPs with DNA (red) show Raman modes characteristic of nucleic acid molecules. Sharp spikes in the signal are cosmic rays, which are removed in standard signal processing. Both the Ag NPs blank and Ag NPs + DNA sample were measured with the same experimental conditions: 30 s exposure time, 10 accumulations.

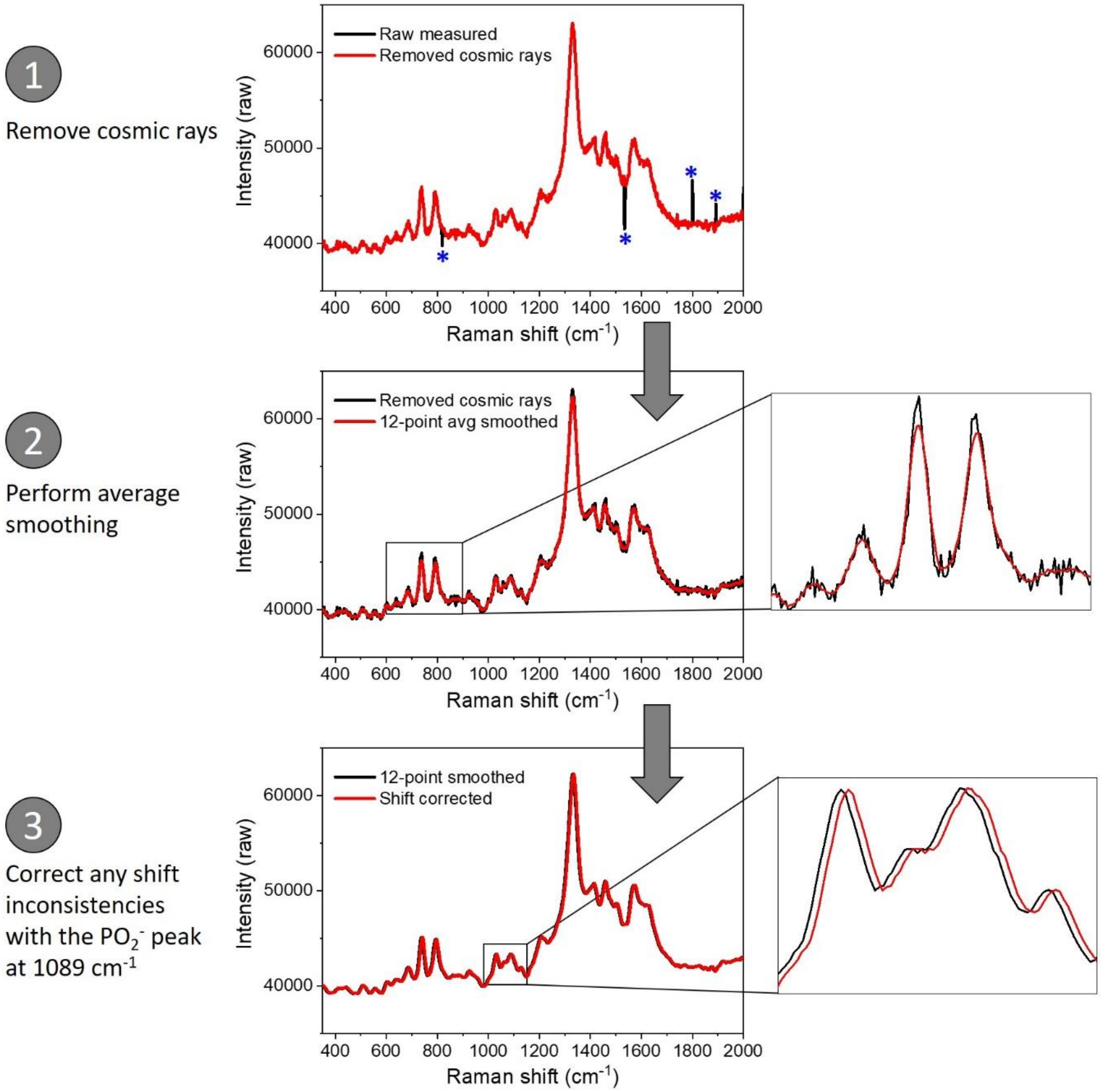


**Figure S2.** SERS signal processing. For optical sequencing analysis, the following processing steps are performed on each spectrum to ensure consistent results: (1) Removing cosmic ray spikes in the spectrum, (2) 12-point average smoothing (±6 in each left/right direction), (3) Correcting small inconsistencies in the 1089 $cm^{-1}$ $PO_2^-$ normalization peak.

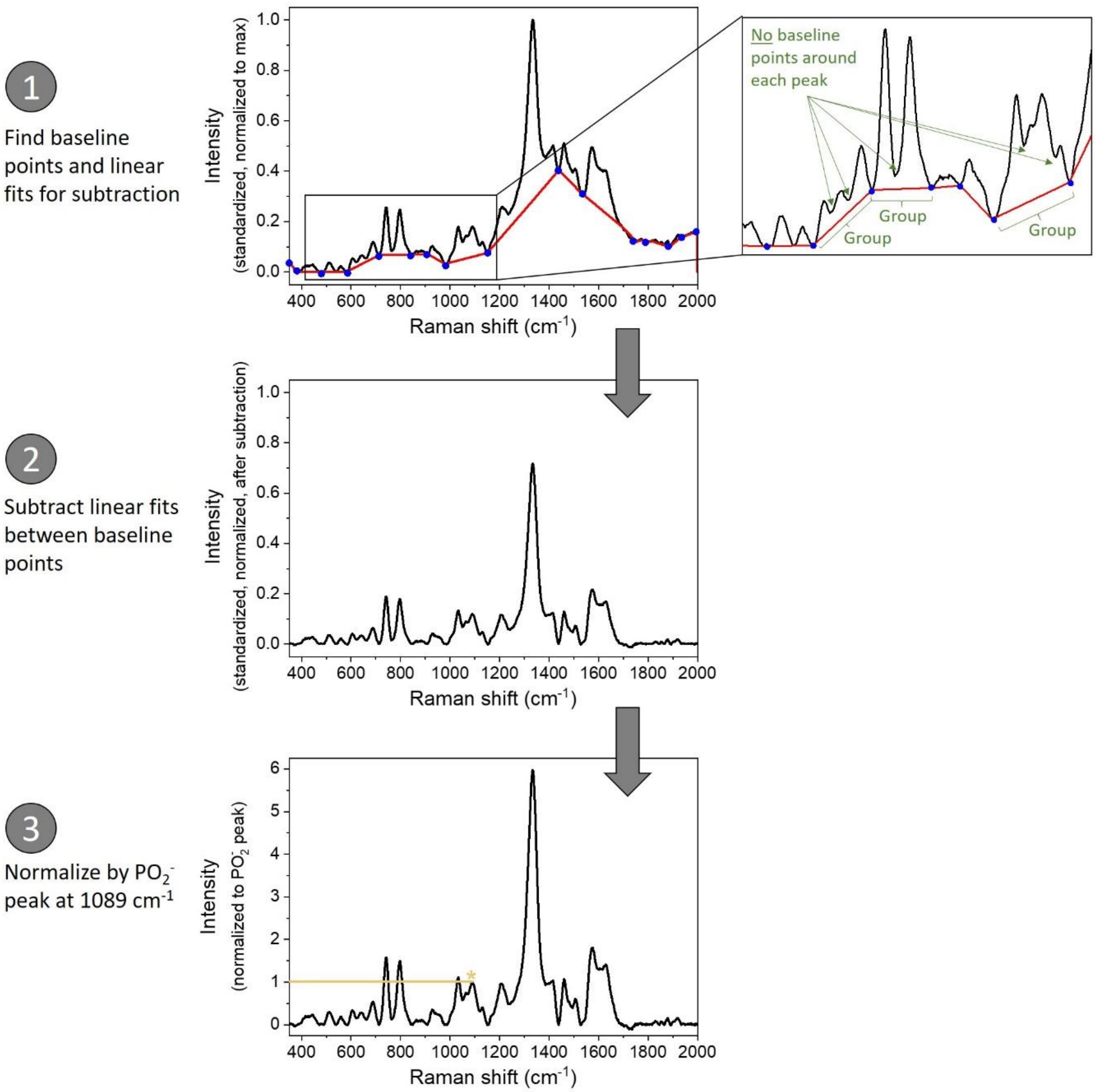


**Figure S3.** SERS signal normalization. For optical sequencing analysis, the following normalization steps are performed on each spectrum to ensure consistent results: (1) Finding all baseline points along the spectrum between which linear fits are constructed – note that the spectrum is first standardized such that the lowest point is brought to zero and normalized by the maximum such that the new maximum is one, (2) Subtracting the linear fits between the baseline points, (3) Normalizing by the 1089 $cm^{-1}$ $PO_2^-$ peak intensity. Shown in the zoomed region of the first step, baseline points cannot simply be set around each peak, but consistently around groups of peaks that give a combined raised signal in the nearby regions.

**Table S1.** Raman peak mode assignments[1–7]

| Raman shift (cm-1) | Assignment |
|---|---|
| 460 | Ring bend (T)* |
| 511 | Ring deformation (mostly G) |
| 600 | Ring bend (C) * |
| 640 | Ring deformation (mostly G) |
| 690 | Ring breathing (G)* |
| 740 | Ring breathing (A)* |
| 790-800 | Ring breathing (C, T) |
| 873 | Backbone vibration, O-P-O stretch |
| 1026 | Deoxyribose + C |
| 1089 | $PO_2^-$ symmetric stretch |
| 1200 | C-H bend |
| 1233 | Ring stretch (C, T) |
| 1295 | Deoxyribose + C, T |
| 1338 | Ring modes (A, G) |
| 1450 | Ring modes (C, T) |
| 1480 | Ring modes (A, G) |
| 1508 | Ring mode (A) |
| 1575 | Ring modes (A, G) |
| 1639 | C=O stretch (mostly C, T) |

*Signature peaks for identifying A-G-C-T content in BOS

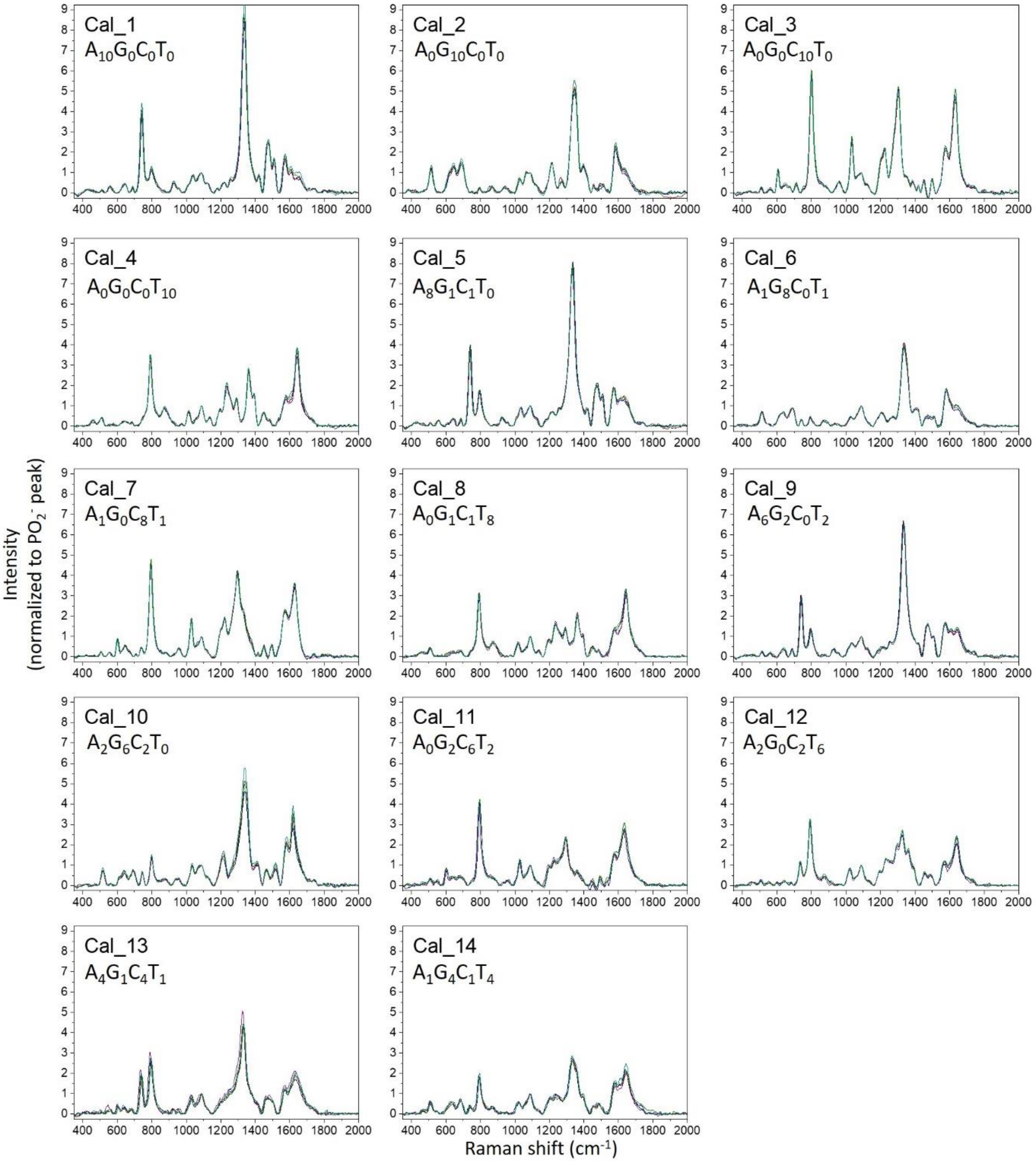


**Figure S4.** All SERS measurements from calibration blocks. Raman spectra are shown with standard signal processing and normalization for each of the 14 calibration 10-mer blocks. Each calibration block was measured with five technical replicates under the same experimental conditions (30 s exposure time, 10 accumulations). Consistency in the measurements is observed from the nearly identical signal from each of the five replicates overlaid together as shown here.

**MEGARes gene**

**Database No.:** 1273

**Class:** Class_D_betalactamases

**Sub-class:** OXA

**Name:** Bla|OXA-50|AM117128|1-789|789|betalactams|Class_D_betalactamases|OXA

**Sequence:**
ATGCGCCCCCTCCTCTTCAGTGCCCTTCTCCTGCTTTCCGGGCATACCCAGGCCAGCGAATGGAATGACAGCCAGG
CCGTGGACAAGCTATTCGGCGCGGCCGGGGTGAAAGGCACCTTCGTCCTCTACGATGTGCAGCGGCAGCGCTATG
TCGGCCATGACCGGGAGCGCGCGGAAACCCGCTTCGTTCCCGCTTCCACCTACAAGGTGGCGAACAGCCTGATCG
GCTTATCCACAGGGGCGGTTAGATCCGCCGACGAGGTTCTTCCCTATGGCGGCAAGCCCCAGCGCTTCAAGGCCT
GGGAGCACGACATGAGCCTGCGCGAGGCGATCAAGGCATCGAACGTACCGGTCTACCAGGAACTGGCGCGGCGC
ATCGGCCTGGAGCGGATGCGCGCCAATGTCTCGCGCCTGGGTTACGGCAACGCGGAAATCGGCCAGGTTGTGGA
TAACTTCTGGTTGGTGGGACCGCTGAAGATCAGCGCGATGGAACAGACCCACTTTCTGCTCCGACTGGCGCAGGG
AGAATTGCCATTCCCCGCCCCGGTGCAGTCCACCGTGCGCGCCATGACCCTGCTGGAAAGCGGCCCGGGCTGGGA
GCTGCACGGCAAGACCGGCTGGTGCTTCGACTGCACGCCGGAACTCGGCTGGTGGGTGGGCTGGGTGAAGCGCA
ACGAGCGGCTCTACGGCTTCGCCCTGAACATCGACATGCCCGGCGGCGAGGCCGACATCGGCAAGCGCGTCGAA
CTGGGCAAGGCCAGTCTCAAGGCTCTCGGGATACTGCCCTGA

**Figure S5.** OXA β-lactamase (class D) gene and blocks. Taken from the MEGARes antibiotic resistance database.[8]

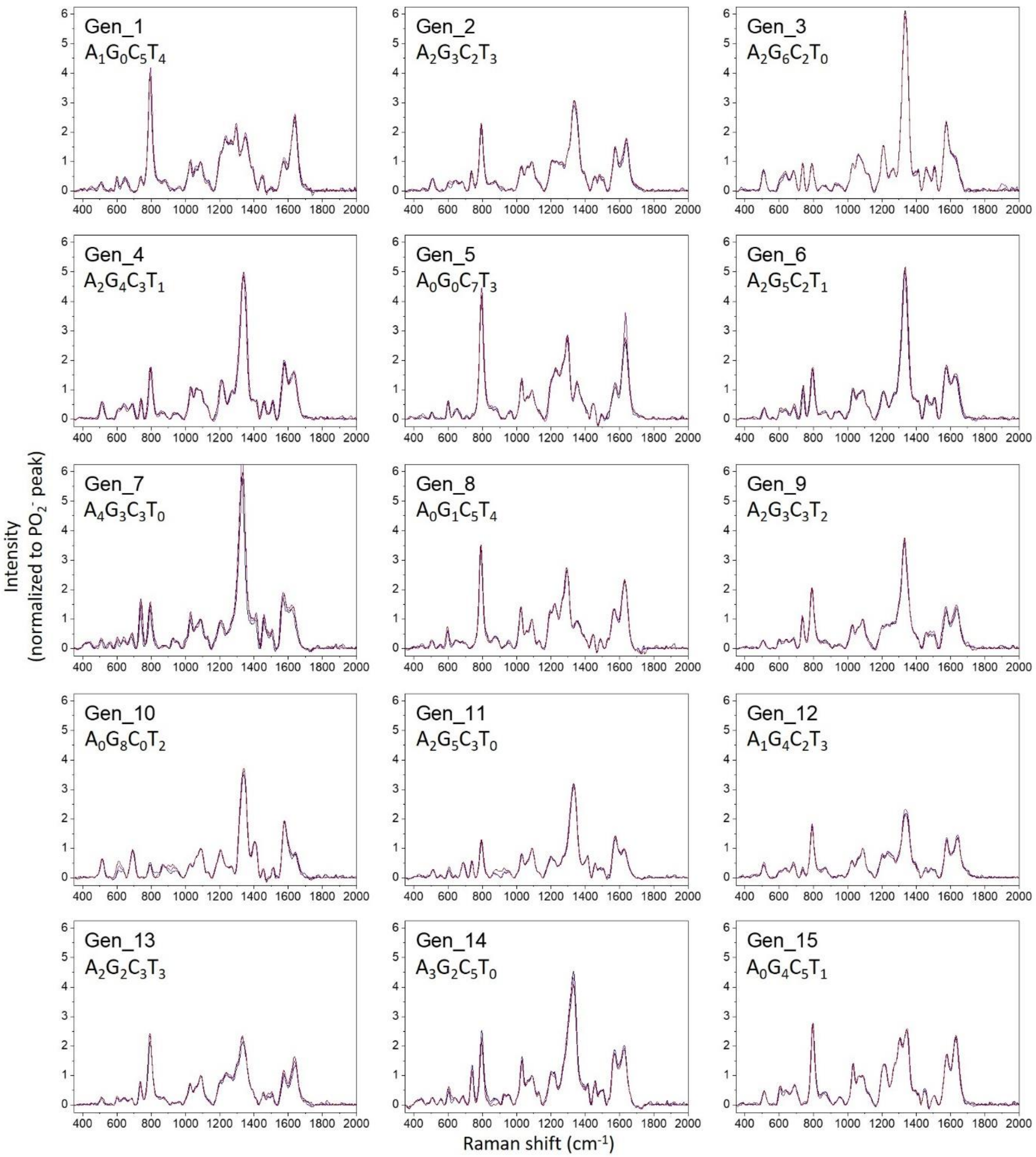


**Figure S6.** All SERS measurements from OXA gene blocks. Raman spectra are shown with standard signal processing and normalization for each of the 15 OXA gene 10-mer blocks. Each calibration block was measured with three technical replicates under the same experimental conditions (30 s exposure time, 10 accumulations). Consistency in the measurements is observed from the nearly identical signal from each of the three replicates overlaid together as shown here.
